# Fractionated radiation alters the extracellular matrix produced by muscle-invasive bladder cancer cells

**DOI:** 10.1101/2025.04.08.647735

**Authors:** C. Guerrero Quiles, S. Fahy, M. Bartak, Gonzalez A. Julia, E. Powel, R. Reed, K. Reeves, A. Baker, Peter Hoskin, Nicholas D. James, Emma Hall, Robert A. Huddart, Nuria Porta, C.M. West, Luisa V. Biolatti, A. Choudhury

## Abstract

Muscle-invasive bladder cancer (MIBC) is a prevalent disease that can be treated with radiotherapy, but has a poor prognosis. Radiation-induced extracellular matrix (ECM) remodelling and fibrosis can induce tumour resistance and recurrence, but has not been studied in MIBC. Here we aimed to characterise the impact of radiation on the ECM composition of MIBC.

**Materials and Methods:** Three MIBC cell lines (T24, UMUC3, J82) were treated with fractionated radiation. We used proteomics to analyse the ECM composition produced by surviving cancer cells and immunofluorescence to investigate changes in the morphology and number of ECM fibres. We evaluated the RNA expression of identified ECM proteins (FN1, COL5A1, COL1A1, TNF6AIP6, FLG) in one cystectomy (TCGA-BLCA, n=397) and two radiotherapy (BC2001, n=313; BCON, n=151) cohorts.

**Results:** There were 613 proteins affected by radiation (p_adj_<0.05, fold change >2 or <-2), 68 of which were ECM-associated proteins. There was a general increase in proteases and protease regulators but heterogeneity across cell lines. Enrichment analysis showed ECM organisation was the primary pathway affected. Immunofluorescence confirmed radiation affected ECM structure, generally, reducing the number, length and width of fibres. High *FN1*, *COL5A1*, *COL1A1, TNF6AIP6* and *FLG* mRNA levels were adverse prognostic markers in clinical cohorts.

**Conclusion:** Radiation alters the composition and structure of the ECM produced by MIBC. As a proof-of-concept, we showed the expression of radiation-affected ECM genes are prognostic markers in MIBC. Future studies should validate these radiation-induced ECM changes in clinical samples.

## 1. Introduction

Bladder cancer is a prevalent disease with >550,000 new cases yearly worldwide which can be classified as non-muscle invasive (NMIBC) or muscle-invasive (MIBC). NMIBC has high 5-year survival rates, >95% for *in situ* and 75% for localised tumours (Halaseh et al., 2022). However, MIBC prognosis is poor, with a 21% 5-year overall survival rate in England (Catto et al., 2023). UK standard-of-care (SOC) treatment for MIBC is radical cystectomy or radiotherapy (55Gy, 2.75Gy daily over 4 weeks), which have similar outcomes (Kimura et al., 2020; Kotwal et al., 2008). However, although radiotherapy allows for bladder preservation, only 30% of MIBC patients in the UK choose this treatment (Kimura et al., 2020; Kotwal et al., 2008), highlighting a need for research to further improve radiotherapy outcomes.

Radiotherapy resistance mechanisms affect recurrence and poor outcomes for MIBC patients who chose radiotherapy over cystectomy. Many mechanisms are associated with radiotherapy resistance, such as DNA damage repair upregulation, cell-cycle arrest and hypoxia *(Wu et al., 2023)*. One less explored mechanism is the development of tumour fibrosis and extracellular matrix (ECM) remodelling. The ECM is a complex and dynamic network of macromolecules which offers mechanical and structural support to surrounding cells. It plays a key role in several functions, including cell proliferation, growth, differentiation and migration (Winkler et al., 2020). It is composed of over 1,000 different proteins, including collagen (COL), laminin (LAM), fibronectin (FN1) and cell-binding glycoproteins (Naba et al., 2012). During tumour progression, there is increased secretion and crosslinking of FN and COL, leading to desmoplasia and tumour fibrosis (Winkler et al., 2020). Fibrosis is a poor prognostic marker in cancer, which promotes tumour progression (Piersma et al., 2020), and evidence suggests it can directly impact radiotherapy resistance. Cordes *et al*. highlighted radiation increases integrin expression, leading to a 10-fold enhancement in adhesion to LAM and FN and increased radioresistance (Cordes et al., 2002). Handschel *et al*. found increased integrin and cell adhesion molecule expression in head & neck cancer patients after radiotherapy (Handschel et al., 1999). More recently, Jin *et al*. showed a direct association with ECM stiffness, with higher stiffness levels promoting radioresistance in cervical cancer cells by altering apoptotic processes (Jin et al., 2022).

The association of fibrosis with radioresistance is especially relevant during radiotherapy, as radiation induces fibrosis (Yu et al., 2023). This relationship is well-acknowledged; myofibroblasts in the tumour-adjacent tissue increase the deposition and crosslinking of ECM components (e.g. COL, FN1), impairing patient outcomes (Yu et al., 2023). For example, Streltsova *et al*. showed radiotherapy alters COL structures within the bladder in a cohort of 105 patients with cervix or endometrial cancer (Streltsova et al., 2021). Therefore, early radiotherapy-induced fibrosis has the potential to drive radioresistance, especially if occurring within the tumour. As a proof-of-concept, Politko *et al*. showed irradiation alters the expression of proteoglycan (versican, decorin) and glycosaminoglycan (brevican) in the ECM of glioblastoma mouse models (Politko et al., 2021). Targeting the ECM during radiotherapy has recently been proposed to improve treatment outcomes (Deng et al., 2024), but the implementation of ECM-targeted therapies is impaired by a lack of comprehensive studies of the ECM produced by cancer cells during radiotherapy. Therefore, we aimed to characterise the ECM composition produced by MIBC cells that survived fractionated radiation partially mimicking SOC (27.5Gy, 2.75Gy daily over 2 weeks).

## 2. Methodology

### 2.1. Cell culture

T24, UMUC3 and J82 cells were acquired from the American Type Culture Collection (ATCC; Virginia, USA). Cells were grown in McCoy’s 5A with L-glutamine (Gibco, Waltham, USA), supplemented with 10% foetal bovine serum (FBS; Sigma-Aldrich, Missouri, USA). Cells were routinely authenticated and tested for mycoplasma. Cells were maintained in a tissue culture incubator (Leec Culture Safe CO2, Appleton Woods, Birmingham, UK) at 37°C and 5% CO_2_.

### 2.2. Irradiation

T24, J82 and UMUC3 cell lines were irradiated using an Xstrahl CIX3 irradiator (Xstrahl, Camberley, UK). The cells received 2.75Gy Monday to Friday at a voltage of 300Kv and a current of 10 mA for 2 weeks (total dose: 27.5Gy). The fractionated schedule was chosen to partially mimic the current UK SOC radiotherapy for MIBC (Choudhury et al., 2021). Non-irradiated control cells were simultaneously maintained and “mock-irradiated”. After irradiation, cells were maintained for four additional weeks in cell culture without any treatment to allow for recovery.

### 2.3. Production of cell-derived ECMs (CDMs)

After recovery, cells were grown for 7 days in normal tissue culture conditions. Cell-derived matrices (CDMs) were extracted as previously described (Humphries et al., 2022). Briefly, decellularisation was performed with a decellularisation buffer (20 mM H_4_OH, 0.5% Triton X-100 in phosphate-buffered saline [PBS]) and CDM was recovered by scraping with 2X SDS buffer (4% (w/v) SDS, 10% (w/v) glycerol, 50 mM Tris HCl, 0.005% (w/v) bromophenol blue, 20% (v/v) mercaptoethanol). To increase protein concentration, CDM samples were acetone precipitated as previously described (Jones et al., 2015). In short, four volumes of acetone were added, and samples were incubated at −80°C overnight. The supernatant was removed, samples washed with the same volume of acetone, and the resulting pellet air-dried at room temperature. Samples were then resuspended in 2X SDS buffer in a thermomixer shaker (1000rpm, 20 min, 70°C). Protein concentration was determined using InstantBlue (Abcam, Cambridge, UK) after SDS-PAGE electrophoresis (45 min, 200 V; 4-12% Bis-Tris gels; Thermo Fisher Scientific), using protein samples of known standard concentrations as previously described (J. Robertson et al., 2017).

### 2.4. Mass spectrometry (MS)

MS analysis was performed following previously described protocols (Humphries et al., 2022; J. Robertson et al., 2015). In short, 5 µg of protein was loaded onto a 4-15% agarose gel (Thermo Fisher Scientific for SDS-PAGE electrophoresis (3 min, 200V; Thermo Fisher Scientific). Protein bands were stained with InstantBlue Coomassie (Abcam), sectioned, and in-gel trypsin digested before being analysed by tandem liquid chromatography-mass spectrometry (LC-MS/MS) using an UltiMate 3000 Rapid Separation LC system (Thermo Fisher Scientific) coupled with an Orbitrap Elite Mass Spectrometer (Thermo Fisher Scientific).

Following previously described protocols (Byron et al., 2015; J. Robertson et al., 2015), MS data was analysed using an in-house Mascot Server (v. 2.5.1; Matrix Sciences) with mass tolerances of 0.4 Da and 0.5 Da for precursor and fragment ions, respectively. Data was validated using Scaffold (v. 4.6.3; Thermo Fisher Scientific) with an identification threshold of 90% at the peptide level, with at least one unique validated peptide (0.1% estimated false discovery rate). Protein identification, normalisation, and fold-change and p-values calculations were performed with Protein Discoverer (v. 2.5.0.400; ThermoFisher Scientific).

### 2.5. Immunofluorescence

Non-previously irradiated cells were single-dose irradiated (8Gy; 300Kv, 10mA) using an Xstrahl CIX3 irradiator (Xstrahl). Non-irradiated control cells were simultaneously “mock-irradiated”. Cells were allowed to recover in a tissue-culture incubator overnight (37°C, 5% CO_2_) and fixed with 8% (w/v) paraformaldehyde solution (10 min, room temperature). Samples were blocked (1 h, room temperature) using blocking buffer (1% [w/v] BSA in PBS) and sequentially stained: (1) overnight incubation (4°C) with rabbit polyclonal antibodies for FN (Sigma-Aldrich, 1:300 dilution), COL5 (Novus Biological, 1:300 dilution), or COL1 (Novus Biological, 1:300 dilution) in blocking buffer; (2) permeabilisation with 0.3% (v/v) Triton-100X in PBS (30 min, room temperature), followed by overnight incubation (4°C) with vinculin (Sigma-Aldrich, 1/300 dilution) and PXN (BD Biosciences, New Jersey, USA; 1/300 dilution) monoclonal mouse antibodies in blocking buffer; (3) incubation with 488 nm Alexa Fluor anti-rabbit (Thermo Fisher Scientific; 1/500 dilution), Alexa Fluor 546 nm anti-mouse (Thermo Fisher Scientific; 1/500 dilution) secondary antibodies, in blocking buffer (2 h, room temperature); (4) nuclei staining (5 min, room temperature) with 300 mM DAPI (Thermo Fisher Scientific) in PBS. Samples were imaged using high-content screening (PerkinElmer Opera Phenix; PerkinElmer) using three confocal spinning disk lasers (405 nm 50mW, 488 nm 50mW, 561 nm 50mW) with a fixed light path system. Three Zyla sCMOS cameras, 2160×2160 pixels, 6.5um pixel size (Andor, Belfast, UK) were set up for each dedicated light path. Twenty fields of view were acquired with a Z range of 34.5µm in 1.5µm steps using the Zeiss W Plan-Apochromat X20 water objective NA 1.0 WD 1.17 mm. Images were analysed using the Harmony software (V. 4.9; PerkinElmer, Massachusetts, USA), reconstructing the acquired field-of-view images as a 2D maximum projection object. Fibres were defined as any object stained with anti-FN, COL1, or COL5 antibodies with roundness <0.8 and an intensity signal >70% of the threshold intensity background signal. DAPI staining was used to estimate the total amount of cells for each acquired image.

### 2.6. Clinical cohorts

Three MIBC cohorts with whole transcriptomic data were used: TCGA-BLCA (n=397), BC2001 (n=313), BCON (n=151). TCGA-BLCA details were previously detailed (A. G. Robertson et al., 2017). Transcriptomic and clinical data is publicly available and was downloaded from the cBioPortal repository (de Bruijn et al., 2023). Patients reported to have had radiotherapy were removed (n=10) and the cohort was considered a cystectomy cohort. BC2001 (NCT00024349; assessing the addition of chemotherapy to radiotherapy) (James et al., 2012) and BCON (NCT00033436; assessing hypoxia-modifying therapy combined with radiotherapy) (Hoskin et al., 2010) are multicentre, randomised, phase 3 trials. Both trials followed local practice radiotherapy (64 Gy in 32 fractions over 6.5 weeks or 55 Gy in 20 fractions over 4 weeks). Whole-transcriptomic methodology and trial details were previously published for both cohorts (Choudhury et al., 2021b; Hoskin et al., 2010; James et al., 2012; L. Yang et al., 2017). For all cohorts, the primary endpoint was 5-year overall survival (OS), defined as the time from the date of randomisation to the date of death up to a 5-year cut-off. Patients were stratified based on the media gene expression for each gene of interest (*FN1*, *COL5A2*, *COL1A1*). Patients were then classified as “low” (<50%) or “high” (≥50%) based on the median expression levels for each gene.

### 2.7. Data analysis

Data filtering, clustering, enrichment, and prognostic analyses were carried out using RStudio (v. 1.5033) using the following packages from The Comprehensive R Archive Network (CRAN, https://cran.r-project.org/): *org.Hs.eg.db*, *AnnotationDbi*, *ClusterProfiler*, *DOSE*, *ggplot2*, *enrichplot, ReactomePA, GoSemSIM, ggVennDiagram*, *tibble*, *survival*, *survminer*. To avoid data biases in the analysis from non-ECM contaminant proteins, data was filtered using the Matrisome Project database as a reference (Shao et al., 2023), excluding all non-ECM proteins from the analyses.

Immunofluorescence data analysis normality was tested using the Shaphiro-Wilks test and statistical differences were determined with a Kruskal-Wallis test with p. values correction. All measurements were normalised based on the total estimated number of cells, and fold changes were calculated using non-irradiated controls as a baseline reference. All significant values were estimated in comparison to the baseline non-irradiated controls. Analyses were performed using GraphPad Prism (v. 9.3.1; GraphPad, Massachusetts, USA).

## Results

### Fractionated radiation alters the extracellular matrix (ECM) composition of MIBC cells

Proteomics identified 1,408 proteins (n=1,357 in T24, n=1,062 in UMUC3, n=912 in J82). A total of 613 unique proteins were identified as altered by radiation across the three cell lines (p_adj_<0.05, fold change > 2 or <-2). In T24, 45 proteins were upregulated and 73 downregulated (Figure 1A); UMUC3 had 121 upregulated and 165 downregulated proteins (Figure 1B); whilst J82 had 253 upregulated and 113 downregulated proteins (Figure 1C). Enrichment analysis confirmed CDM samples were significantly enriched in proteins associated with the ECM-related cellular compartment terms including “cell-substrate junction” (14.1% of proteins) and “focal adhesion” (14.2% of proteins). Enriched terms related to non-ECM proteins such as ribosomal and cytosolic subunit were also identified at a lower percentage than those associated with the ECM. Proteomic analysis confirmed that radiation alters ECM composition produced by bladder cancer cells.

**Figure 1:**
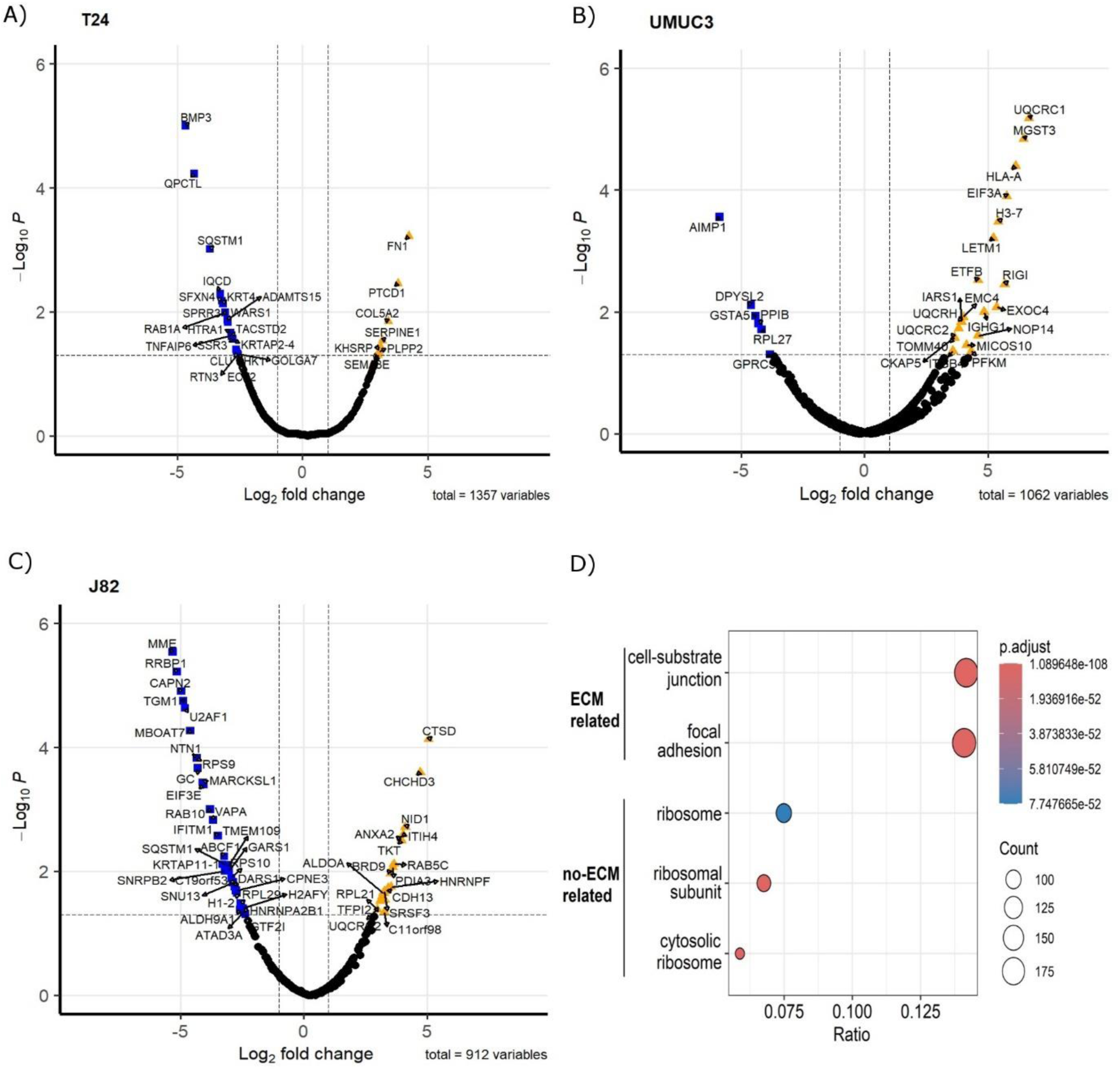
Fractionated radiotherapy partially mimicking SOC (27.5Gy, 2.75Gy daily over 2 weeks) alters the protein composition of the ECM produced by muscle-invasive bladder cancer cell lines. Volcano plots represent significantly upregulated (fold change >2, p.adj. <0.05) and downregulated (fold change <-2, p.adj. <0.05) proteins in cell-derived extracellular matrices (CDMs) for T24 (A), UMUC3 (B) and J82 (C) cell lines. Cellular compartment GO Term enrichment analysis (D) confirms CDMs protein fractions are enriched in ECM-liked proteins. Upregulated proteins are represented as orange triangles; downregulated proteins are represented as blue squares. Three biological repeats were analysed per cell line.

### Fractionated radiation mainly affects the expression of ECM non-structural proteins

We used the Matrisome database as a reference to remove any non-ECM proteins from the generated datasets. After filtering, we identified with high confidence a total of 68 ECM proteins affected by radiation (Figure 2A). Non-structural ECM proteins were the most prevalent, which included growth factors (e.g. platelet-derived growth factor [PDGF] B, fibroblast growth factor [FGF] 5), cytokines (e.g. chemokine CXC motif ligand [CXCL] 1, CXCL2), proteases (e.g. adamalysin [ADAM] TS12, ADAMTS19) and proteases-regulators (e.g. serpin [SERPIN] E1, SERPINF2) (Figure 2A). However, we also observed changes in ECM structural proteins (e.g. COL5A2, FN1). Radiation-induced changes in the ECM composition were cell-line dependent, with only two significant in common across all three cell lines (Filaggrin [FLG], tumour necrosis factor-alpha induced-protein 6 [TNFAIP6]) (Figure 2B). Of interest, although COL upregulation was observed (e.g. COL5A2 in T24), several collagen types (e.g. COL4A3 in UMUC3, COL7A1 in J82) were downregulated. Pathway enrichment analysis confirmed radiation mostly alters the “ECM organisation” affecting >38% of all significant proteins (Figure 2C). Interestingly, Molecular function enrichment highlighted “peptidase regulators” (19.4%) and “growth factors” (17.9%) as the most common types of radiation-altered proteins, with “ECM structural components” (16.4%) ranking third (Figure 2D). To confirm these results, we performed another enrichment analysis using the Matrisome database ontology terms designed specifically for ECM analysis. The analysis validated the previous results, with 76% of significant proteins being “ECM-associated”, whilst the “Core ECM” structure only comprised 24% of proteins (Figure 2E). A deeper analysis confirmed that “secreted factors” (37%) and “ECM regulators” (23%) were the most affected protein types following fractionated irradiation. “ECM-affiliated proteins” (13%) was the third most affected “ECM associated” protein type. Regarding the “Core ECM” proteins, “glycoproteins” (19%) were the main affected type, with only 6% and 1% of significant proteins being “collagens” and “proteoglycans”, respectively. GO Molecular function enrichment validated these results, showing a consistent enrichment in growth factors, peptidase, and peptidase regulators (Figure S1a). A general increase in peptidase and peptidase regulators was observed (Figure S1b-d). In addition, a significant enrichment in cytokines and glycosaminoglycan binding molecules was also observed (Figure S1). Overall, our analysis suggests fractionated radiation mostly affects the ECM organisation indirectly by altering the expression of ECM regulators rather than directly altering the expression of ECM structural proteins.

**Figure 2:**
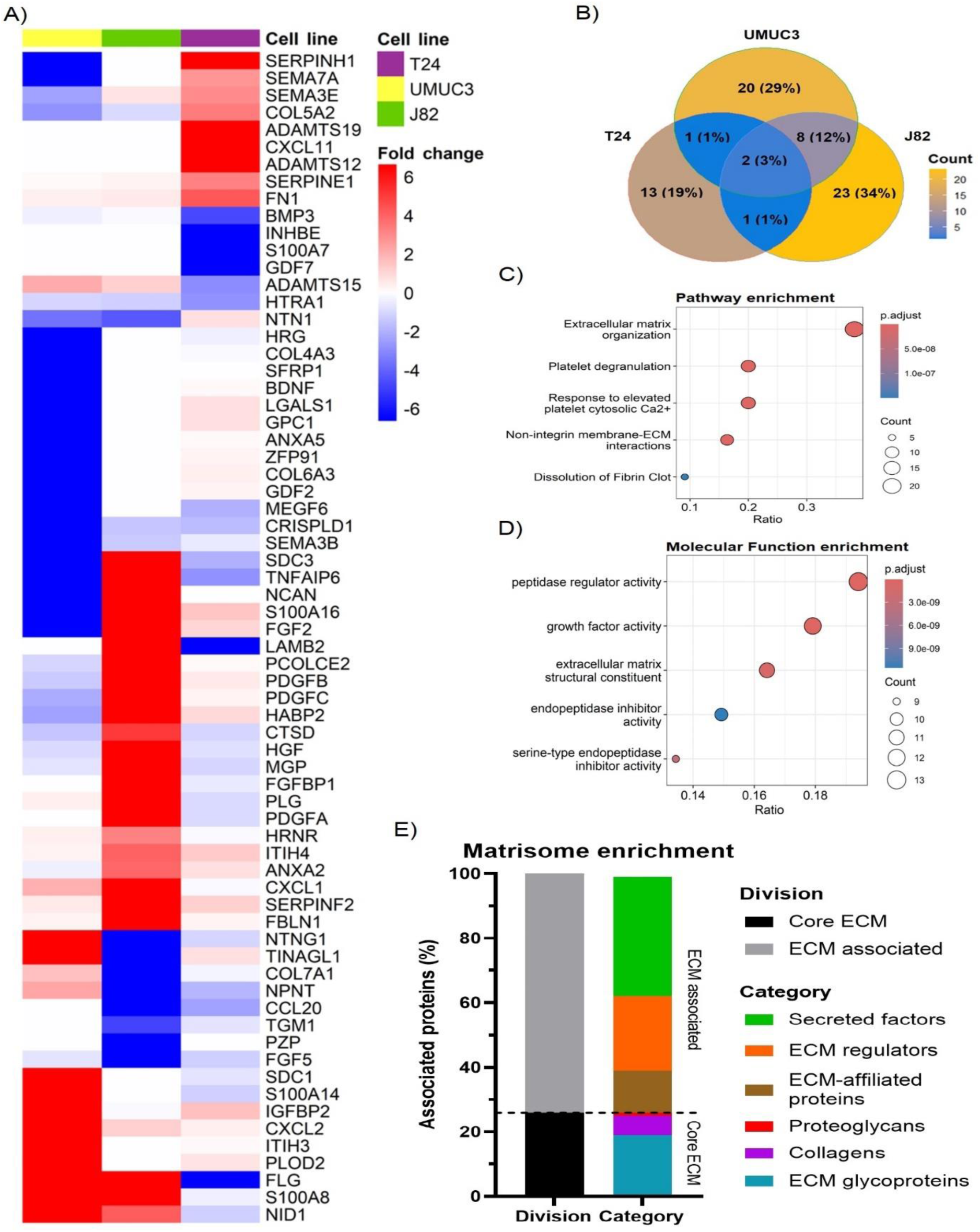
Fractionated radiotherapy partially mimicking SOC (27.5Gy, 2.75Gy daily over 2 weeks) alters the expression of 68 extracellular matrix (ECM) proteins, which are mostly associated with “peptidase regulator activity” and “ECM organisation”. Heatmap (A) represents the fold change for all identified ECM proteins significantly upregulated (fold change >2, p.adj. <0.05) and downregulated (fold change <-2, p.adj. <0.05) in T24, UMUC3 and/or J82 cell lines. Venn diagram (B) shows the overlapping among the significantly up and downregulated proteins identified in each cell line, highlighting cell line variability. Reactome pathway enrichment (C) shows radiotherapy mostly alters the ECM organisation. GO Term Molecular Function enrichment analysis (D) shows radiotherapy mostly affects peptidase regulators, followed by growth factors and ECM structural components. Ratio (B, C) represents the % of total proteins associated with each term (0 – 1 scale). Matrisome terms enrichment (E) shows >70% of altered proteins are ECM-associated factors and protein regulators.

### Fractionated radiation induces diverse ECM regulator mechanisms across cell lines

As the previous analysis highlighted high cell line variability, we performed individual gene enrichment analyses for each cell line to find commonly affected pathways. In all cases, the pathway enrichment analysis highlighted “ECM organisation” as the most affected pathway, linked to 40% of proteins in T24, 37.5% in UMUC3, and 42.9% in J82 (Figure 3A). “ECM proteoglycans”, “non-integrin membrane/ECM interactions”, as well as several beta galactosyl and glycosyltransferase pathways (e.g. B3GALT, B3GAT) were enriched in all cell lines, validating our previous results (Figure 3A). However, the pathway alterations are predicted to be induced by different proteins in each cell line. For example, SERPINs protein family signalling is predicted to regulate the “ECM organisation” in T24 and UMUC3 cell lines (Figure 3B,C), whilst platelet-derived growth factors are predicted regulators only in J82 (Figure 3D). Similarly, beta galactosyl and glycosyltransferase activity is regulated by ADAMs) in T24 (Figure 3B), but by syndecans (SDCs) in UMUC3 (Figure 3C). Our results suggest that radiation affects different ECM regulator mechanisms, which converge by affecting the same pathways.

**Figure 3:**
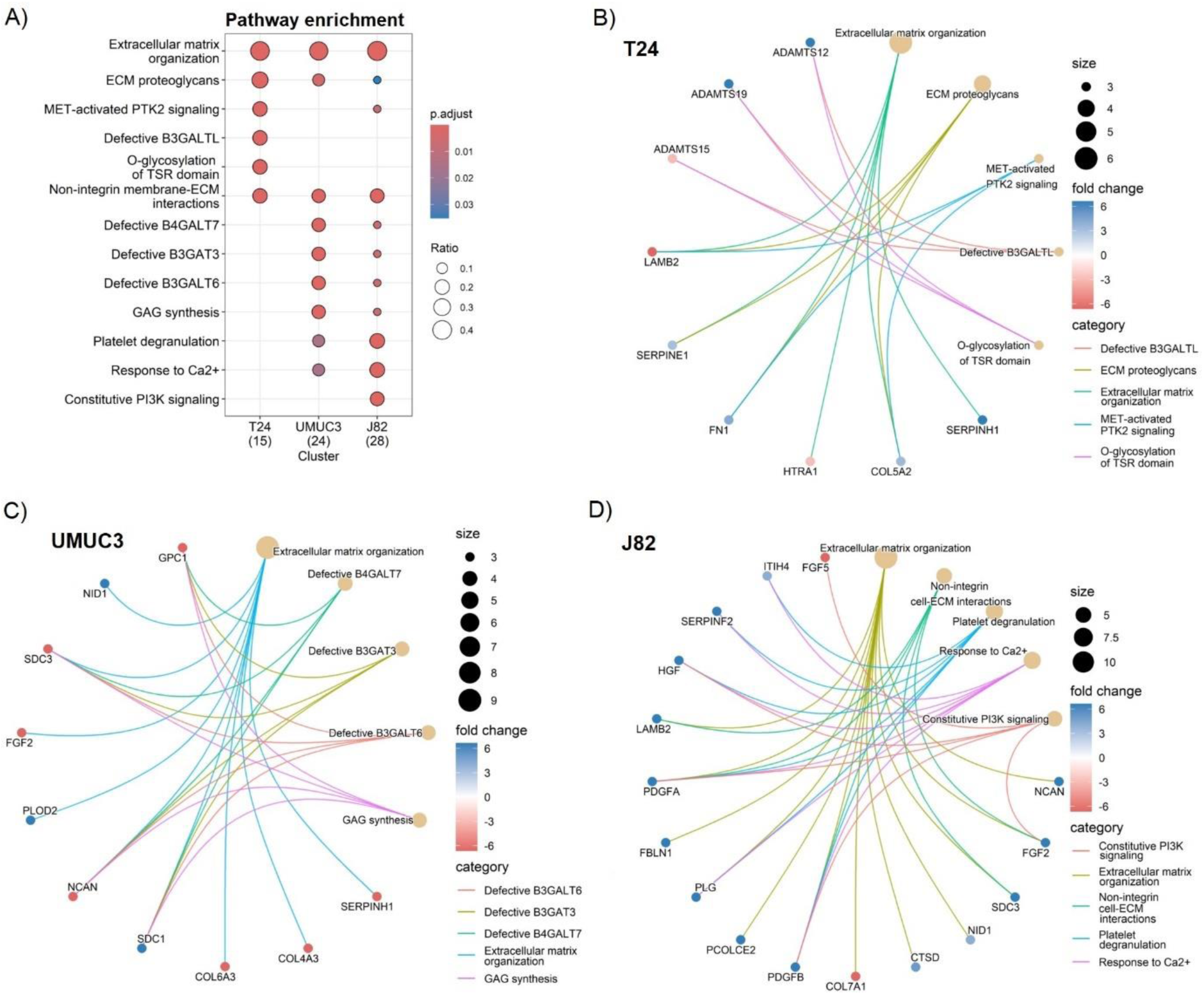
Fractionated radiotherapy partially mimicking SOC (27.5Gy, 2.75Gy daily over 2 weeks) consistently alters the extracellular matrix (ECM) organisation, ECM proteoglycans composition, and the non-integrin cell/ECM interactions across all cell lines. Comparative Reactome pathway enrichment analysis (A) of significantly up (fold change >2, p.adj. <0.05) and downregulated (fold change <-2, p.adj. <0.05) ECM proteins for each individual cell line (T24, UMUC3, J82). Cnetplots show specific associations among significantly up and downregulated proteins and their corresponding enrichment terms for T24 (B), UMUC3 (C) and J82 (C). Ratio (A) represents the % of total proteins associated with each term (0 – 1 scale). A total of n=3 biological repeats were analysed per cell line.

### Irradiation alters structural ECM fibre number and morphology

As ECM organisation was the main affected pathway following irradiation, we studied how irradiation alters the number and morphology of ECM structural proteins that produce fibres (fibronectin [FN1], collagen 5 [COL5], collagen 1 [COL1]). Irradiation generally decreased FN1 (T24, J82), COL5 (T24, J82) and COL1 (UMUC3) fibre numbers in all cell lines (Figure 4A,B). Only T24 showed a significant 10-fold increase in the number of COL1 fibres (Figure 4A,B). Regarding morphology, the effect on fibre length was cell-line dependent; radiation increased fibre length in T24 (FN1, COL1), but reduced fibre length (COL5, COL1) in UMUC3 (Figure 4A,C). The most consistent effect was a decreased COL5 (T24, UMUC3, J82) and COL1 (UMUC3, J82) fibre width. There were no changes in the width of FN1 fibres (Figure 4A,D). Of note, we did not detect FN1 synthesis in UMUC3 cells. Altogether, these results show irradiation alters ECM fibre numbers and morphology, suggesting it affects ECM organisation.

**Figure 4:**
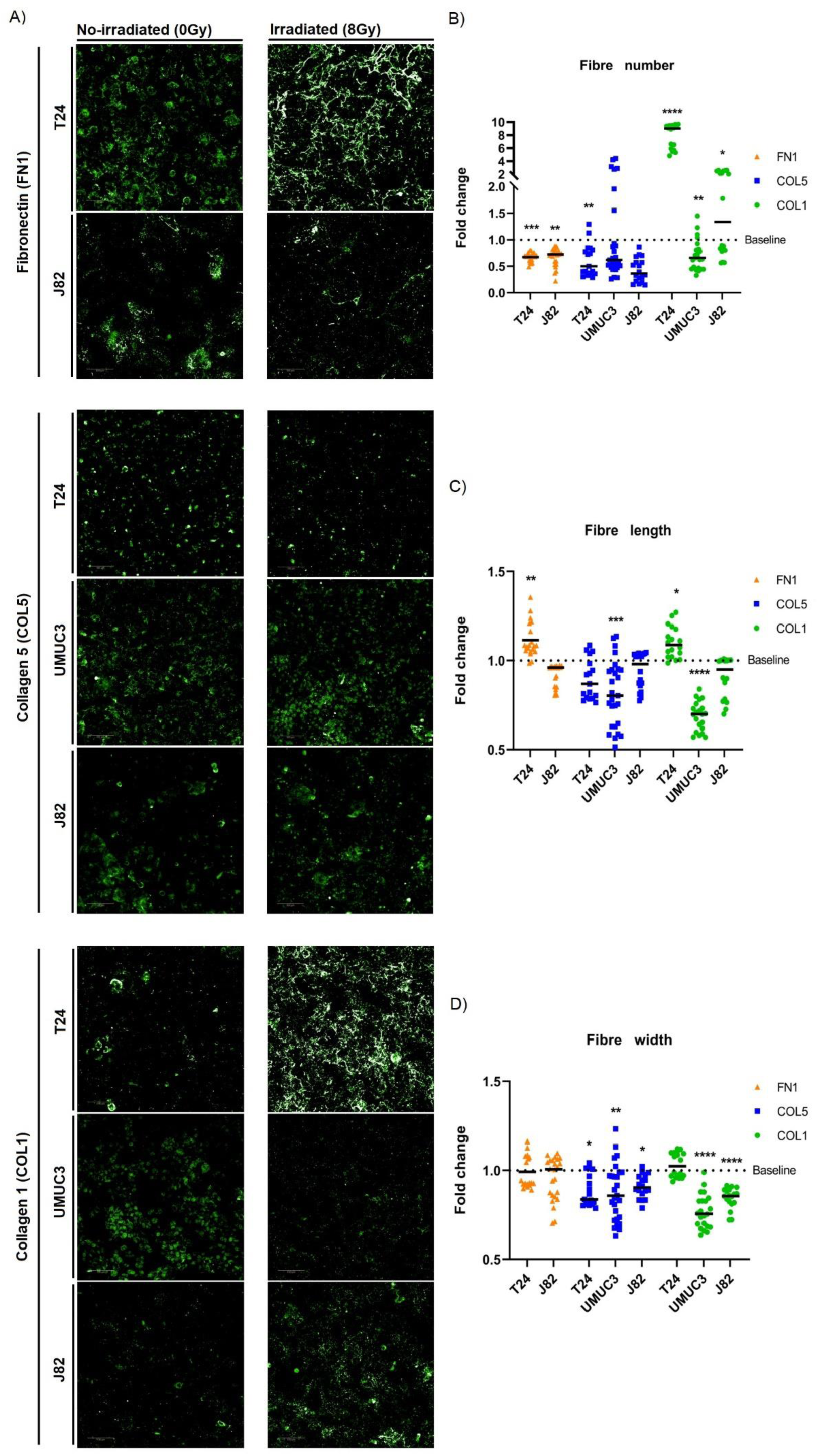
Single-dose radiation (8Gy) affects the morphology and number of ECM fibres. Immunofluorescence staining (A) of fibronectin (FN1), collagen 5 (COL5) and collagen 1 (COL1) fibres for T24, UMUC3 and J82 cell lines. Irradiation consistently decreased fibres numbers (B) for FN1 (T24, J82), COL5 (T24, J82) and COL1 (UMUC3), except for COL1 in T24 (significantly increased). Fibres length (C) alterations were cell-line dependent, with T24 showing a significant increase for FN1 and COL1, whilst UMUC3 had a significant decrease for COL5 and COL1. Fibres width (D) consistently had a significant decreased after irradiation for COL5 (T24, UMUC3, J82) and COL1 (UMUC3, J82) fibres. Significance is defined as p≤0.05, with * for p≤0.05, ** for p≤0.01, *** for p≤0.001 and **** for p≤0.0001.

### *FN1*, *COL5A2* and *COL1A1* expression correlate in MIBC

As we observed a general reduction in *FN1*, *COL5A2* and *COL1A1* fibres after irradiation, we assessed whether their expression levels correlated in MIBC. Figure 5 shows that the expression of the three genes significantly correlated in all three cohorts (p<0.001). FN1 and COL1A1 expression showed the highest consistency in correlation strength, with r>0.8 in TCGA-BLCA (Figure 5C), r>0.5 in BC2001 (Figure 5F), and r>0.7 in BCON (Figure 5I). However, correlation strength variated across cohorts. The range was highest in TCGA-BLCA (r=0.84-0.91), lower in BCON (r=0.62-0.74) and lowest in BC2001 (r=0.34-0.51). Overall, our data shows FN1, COL5A2 and COL1A1 expression correlates in MIBC, although the association may depend on the patient’s pathological characteristics.

**Figure 5:**
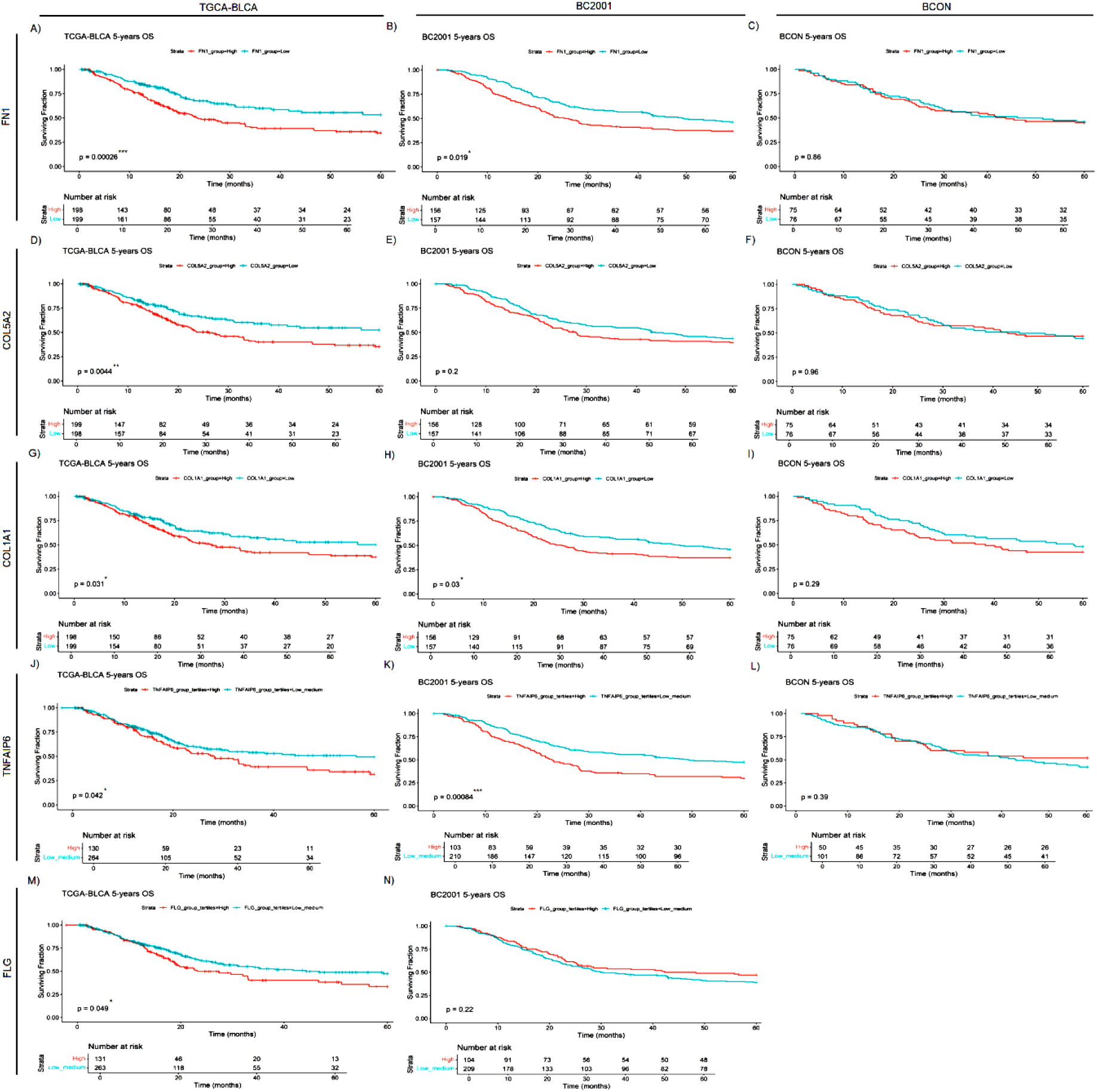
*FN1*, *COL5A2*, *COL1A1, TNFAIP6 and FLG* mRNA expression levels are general prognostic markers for muscle-invasive bladder cancer (MIBC). *FN1* (A-C), *COL5A2* (D-F), *COL1A1* (G-I), *TNFAIP6* (J-L) and *FLG* (M,N) expression was retrospectively validated in one cystectomy (TCGA-BLCA, n=397) and two radiotherapy (BCON [n=151]; BC2001 [n=313]) cohorts. *FN1* expression had significant prognosis in the TCGA-BLCA (A), and BC2001 (B) cohorts. *COL5A2* expression was only significantly prognostic in the TCGA-BLCA (D) cohort. *COL1A1* expression had a significant prognosis in both TCGA-BLCA (G) and BC2001 (H) cohorts. *TNFAIP6* expression was significantly prognostic in the TCGA-BLCA (J) and BC2001 (K) cohorts. Finally, *FLG* expression was only prognostic in the TCGA-BLCA cohort. No *FLG* expression data was available for the BCON cohort. Patients were classified into “High” and “Low” based on each cohort’s median gene expression levels for *FN1* (A-C), *COL5A2* (D-F) and *COL1A1* (G-I) analyses. Patients were classified into “High”, “Medium” or “Low” based on each cohort’s tertiles median gene expression for *TNFAIP6* (J-L) and *FLG* (M,N) expression. Significance was defined as p≤0.05, with * for p≤0.05, ** for p≤0.01, *** for p≤0.001 and **** for p≤0.0001.

### *FN1*, *COL5A2*, *COL1A1*, *TNFAIP6* and *FLG* are prognostic markers in MIBC

Finally, as a proof-of-concept of the clinical relevance of the ECM proteins identified, we analysed the prognostic capacity of *FN1*, *COL5A2*, *COL1A1, TNFAIP6* and *FLG* expression. High *FN1* expression was associated with a poor prognosis in both TCGA-BLCA (p<0.0001; Figure 6A) and BC2001 (p<0.05; Figure 6B). *COL5A2* expression was only prognostic in TCGA-BLCA (p<0.001; Figure 6D). However, *COL1A1* expression was prognostic in both TCGA-BLCA (p<0.05; Figure 6G) and BC2001 (p<0.05; Figure 6I) cohorts. No significance was found for *TNFAIP6* and *FLG* expression after a 50% cohort split into “High” and “Low” groups. However, univariate Cox hazard-risk analysis showed a linear relationship between *TNFAIP6* and *FLG* expression with mortality risk (Figure 2S). After a tertile stratification into “High” (>67%), “Medium” (67–33%) and “Low” (<33%) expression groups, we identified patients with highly induced *TNFAIP6* and *FLG* expression as a subgroup of patients with a potential increase in mortality risk (Figure 2S). “High” *TNFAIP6* expression is a poor prognosis marker when compared to “low and medium” *TNFAIP6* expression levels in the TCGA-BLCA (Figure 6J) and BC2001 (Figure 6K) cohorts. Similarly, “high” FLG expression was also significantly prognostic in the TCGA-BLCA (Figure 5M) cohort when compared to “low and medium” expression levels. None of the studied genes had prognostic value for any analysis in the BCON cohort.

**Figure 6:**
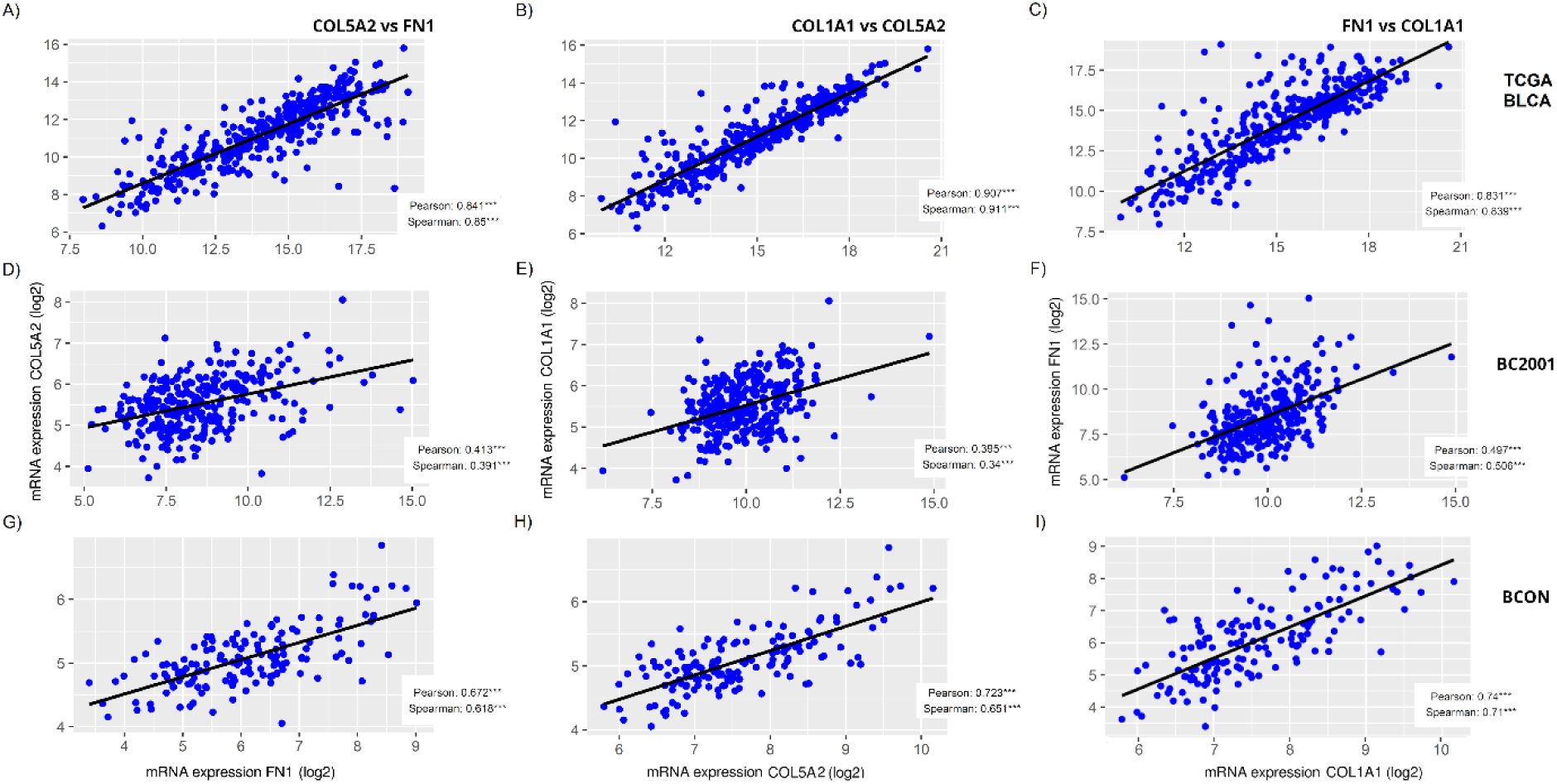
*FN1*, *COL5A2* and *COL1A1* expression correlates in muscle-invasive bladder cancer cohorts. The correlation among *FN1*, *COL5A2* and *COL1A1* mRNA expressions was measured for the TCGA-BLCA (A-C), BC2001 (D-F) and BCON (G-I) cohorts, showing significant a significant correlation in all cases. Both Pearson and Spearman correlation was measured. Significance is represented as * for p<0.05, ** for p<0.01 and *** for p<0.001.

To address whether this fact was associated with half of the BCON cohort patients receiving hypoxia-modifying treatment, we repeated the analyses after splitting the cohort into the two trial arms (Figure S3). *COL1A1* expression was prognostic in the non-CON treatment arm (Figure S3E), suggesting that targeting hypoxia affects the prognostic capacity of ECM genes. Finally, we validated *FN1*, *COL5A2*, *COL1A1, TNFAIP6* and *FLG* expression as independent prognostic markers through a Cox multivariate analysis which included age, gender, T stage and grade as clinical variables. Other added variables were chemotherapy for BC2001 and CON treatment for BCON multivariate analyses. Only *FLG* and *TNFAIP6* had independent prognostic capacity in the TCGA-BLCA (Figure S4A) and BC2001 (Figure S4B) cohorts, with a significant increase of 85% and 48% in mortality risk, respectively. Overall, our results show that the mRNA expression of the ECM proteins identified as affected by radiation is prognostic irrespective of treatment with surgery or radiotherapy.

## Discussion

Here, we used a protein-fractionation approach to characterise the ECM produced by cancer cells. We found our protein fractionation approach led to an enrichment of ECM-related proteins. Eleven per cent of the significant proteins identified (68 of 613) were ECM proteins, a proportion similar to other reports. For example, Naba *et al*. found 10-30% tissue-specific ECM proteins when characterising CDMs from lung and colon tissues (Naba et al., 2012). Similar proportions were identified in other studies as reviewed by Byron *et al* (Byron et al., 2013). Our analysis fits within the expected range and confirms the validity of the approach used.

We found high cell line variability in the ECM proteins affected by radiation, with only two significant in all cell lines (FLG, TNFAIP6). This variability can be associated with the distinctive genomic profile of each cell line. Mutations in ECM genes such as *COL* are common and directly influence the ECM structure (Xu et al., 2019). Furthermore, mutations in master oncogenic regulators like *TP53* also regulate the ECM composition (Kenny et al., 2017; Pickup et al., 2014; Xu et al., 2019). Regarding the cell lines used in this study, no mutations in COL genes are reported for J82, whilst T24 (*COL5, COL6, COL18*) and UMUC3 (*COL4, COL12, COL23*, *COL28*) have distinct mutational profiles. Similarly, T24 cells have a non-sense mutation in *TP53*, while UMUC3 and J82 have miss-sense mutations. As *TP53* regulates the ECM, and is not mutated in T24, it could explain its distinct ECM profile, sharing only 4% of its proteins with UMUC3 and J82, while UMUC3 and J82 shared up to 15%. Regarding FLG and TNFAIP6, no direct previous association with radiotherapy has been reported. Bai *et al*. found that FLG expression regulates PI3K/AKT/mTOR-mediated drug-resistance mechanisms, with FLG expression correlating with drug sensitivity in head and neck cancer (Bai et al., 2021). The PI3K/AKT/mTOR pathway is well-known to induce both drug and radiotherapy resistance mechanisms (Rascio et al., 2021). Therefore, the radiation-induced changes in FLG expression changes observed here might contribute to ECM-induced radioresistance mechanisms. On the other hand, TNFAIP6 is a protein induced by TNF signalling (Guo & Yuan, 2020). TNF signalling has already been reported to regulate radiosensitivity in several cancer cell lines (Dennis et al., 1990). However, TNFAIP6 signalling has barely been explored. A recent pan-cancer meta-analysis showed TNFAIP6 signalling positively correlated with neutrophil infiltration, and was an adverse prognostic factor in multiple cancers (Y. Yang et al., 2024). The positive association of TNFAIP6 with neutrophil infiltration suggests TNFAIP6 expression might associate with a radiation-induced inflammatory response (Franzolin & Tamagnone, 2019; Xia et al., 2018; Zhou et al., 2023). Furthermore, increased neutrophil infiltration promotes radioresistance in sarcoma (Wisdom et al., 2019). However, further validation through radiosensitivity assays is required to confirm whether FLG and TNFAIP6 affect the radiosensitivity of MIBC.

From a broader perspective, our analyses predicted radiation-induced changes in the ECM organisation, with results consistent with previous reports (Yu et al., 2023). Specifically, we found that more than one-third of the ECM proteins affected by radiation were structural. COLs, typically associated with fibrosis and desmoplasia (Winkler et al., 2020), were minimally affected, accounting for only 6% of significant ECM proteins. Notably, the expression of glycoproteins (e.g., FN1) was highly impacted by radiation (19% of the significant ECM proteins). Politko *et al*. showed irradiation increases proteoglycans and glycoprotein expression in glioblastoma, supporting our findings (Politko et al., 2021). Furthermore, Tian *et al*. showed cancer cells and fibroblasts produce distinct ECM profiles. While fibroblast ECMs contain 80-90% of COLs, cancer-cell ECMs have only 5-20% of COLs (Tian et al., 2019). Therefore, due to the limited capacity of cancer cells to produce COLs, proteoglycans likely have a more relevant role in any radiation-induced changes of ECM mechanical properties (e.g. stiffness). This concept is supported by our cell-line independent enrichment analysis, which predicted glycosaminoglycan synthesis, galactosyl and glucuronyl transferases as affected pathways. Post-translational modifications (e.g. glycosylation) are an important step of glycoprotein synthesis (Adams, 2023; Reily et al., 2019), providing context for the radiation-induced changes in glycoprotein levels.

We found ECM regulators and growth factors (e.g. SERPINE1, FGF2, CXCL11) comprised two-thirds of the ECM significant proteins. From an immune perspective, the observed increase in expression of semaphorin (SEMA), S100A proteins, and cytokines (CXCL) suggests the activation of an inflammatory response (Franzolin & Tamagnone, 2019; Xia et al., 2018; Zhou et al., 2023), which is a well-described consequence of radiotherapy within the tumour microenvironment (McKelvey et al., 2018; McLaughlin et al., 2020). Regarding the ECM organisation, ECM fibre crosslinking and organisation are critical regulators of ECM mechanical properties (Winkler et al., 2020). Here, we found increased expression of proteases from the ADAMs family and protease regulators from the SERPINs family in MIBC cells surviving fractionated irradiation. There is also previous evidence associating ADAMs and SERPINs with radiotherapy resistance. SERPIN overexpression induces radioresistance in lung cancer (J. Zhang et al., 2022), and is a poor prognostic marker in bladder cancer (Chuang et al., 2021), whilst ADAMs are upregulated after radiotherapy (Waller & Pruschy, 2021).

Both ADAM and SERPIN families regulate proteolysis of structural ECM proteins (e.g. COL, FN1) (Bonnans et al., 2014; Cox, 2021), suggesting alterations in the ECM fibres’ number and morphology. Indeed, our immunofluorescence analyses showed a general decrease in ECM fibre (FN1, COL5, COL1) numbers, width and length. These results contradict the current knowledge of radiation being an inducer of fibrosis (Yu et al., 2023), as ECM fibre crosslinking is required for fibrosis (Winkler et al., 2020). However, other reports showed similar contradictions. Strelstova *et al*. showed radiotherapy disorganised COL fibres in healthy bladder tissue (Streltsova et al., 2021), whilst increased COL fibres organisation is a key inductor of ECM stiffness and fibrosis (Gilkes et al., 2015; Winkler et al., 2020). It is therefore possible radiotherapy may not always induce fibrosis, which may depend on additional variables (e.g. genomic background). For example, here we observed a 10-fold increase in COL1 fibre number in T24, which supports radiation induces fibrosis. As discussed previously, T24 have a unique mutagenic profile, including a *TP53* non-sense mutations. Therefore, it is possible that specific genomic mutations may be required for cancer cells to induce COL1-mediated fibrosis. In addition, radiation-induced fibrosis has been mainly associated with fibroblasts and myofibroblasts (Streltsova et al., 2021), which produce rich COL-containing ECMs (Tian et al., 2019). It is also plausible that radiation-induced fibrosis depends on the number of fibroblasts in the irradiated tissue. Both these concepts are relevant, as radiotherapy would induce distinct tumour microenvironments depending on the proportion of fibroblasts and the genomic background of the cancer cells. Future genomics analysis in radiobiology should aim to link specific genomic mutations with the capacity to promote ECM fibrogenesis. In addition, alterations in the ECM produced by fibroblasts after fractionated irradiation should also be investigated, following similar approaches to the ones performed in this study.

We also found a significant correlation between the expression of *FN1*, *COL5A2* and *COL1A1* in clinical cohorts, suggesting all three genes behave similarly. Indeed, the expression of all three genes is commonly associated with ECM remodelling and fibrosis (Winkler et al., 2020). Furthermore, high expression of structural ECM genes may be associated with desmoplasia and ECM remodelling, an adverse prognosis factor in several cancer types (Piersma et al., 2020; Quintela-Fandino et al., 2024; Wang et al., 2021). Similarly, we found high *FN1*, *COL5A2*, and *COL1A1* expression associated with a poor prognosis in MIBC. *TNFAIP6* and *FLG* were also prognostic. The association with a poor prognosis was seen in both the cystectomy (TCGA-BLCA) and radiotherapy (BC2001) cohorts, suggesting that high expression of ECM genes associates with a poor prognosis in MIBC irrespective of treatment modality. Our findings agree with those from others as high *COL10A1* expression was previously associated with a poor prognosis in five bladder cancer cohorts (Tian et al., 2019). Furthermore, Chen *et al*. found patients with mutant *FLG* and a sub-group with wild-type *FLG* have a poor prognosis (Chen et al., 2022). These reports support our results, which suggested a subgroup of patients with high *FLG* expression conferred a poor prognosis. Similarly, *TNFAIP6* expression has been reported as an independent adverse prognosis marker in the GSE32894 bladder cancer cohort (Chan et al., 2019). Similar results were found in gastric cancer (X. Zhang et al., 2021).

In multivariable analysis only *FN1* and *TNFAIP6* retained independent prognostic significance. The fact that *COL5A2* and *COL1A1* were not validated as independent prognostic markers is likely due to their high correlation with *FN1*, suggesting they provide no additional information. Interestingly, we found *COL1A1* was only prognostic in BCON patients not receiving hypoxia-modifying CON treatment. Hypoxia has been reported to promote fibrosis and ECM stiffness (Gilkes et al., 2015). Therefore, CON treatment may reduce the impact ECM has on patient survival.

Our study has several limitations. We only studied the effect of fractionated irradiation in cancer cells, as we had no fibroblasts available. We neither studied protein expression nor fibre numbers in the clinical cohorts and only had access to data derived from diagnostic biopsies.

The prognostic capacity of the genes reported here may be due to a general fibrotic process, which is well-associated with tumour progression (Pickup et al., 2014). Our *in vitro* data suggests radiation can decrease ECM fibre number, hence impairing the fibrotic process (Winkler et al., 2020). To address these apparent contradictions and clinically validate radiotherapy-induced ECM changes we would need to access tumour samples taken during radiotherapy (probably impossible). However, investigating ECM alterations in fibroblasts after fractionated irradiation is feasible and should be attempted in future studies, as it would provide context to clarify how fibrosis is induced by irradiation. Future studies should also evaluate the prognostic capacity of COL and FN1 fibres and expression levels in radiotherapy recurrence samples, to confirm the prognostic effect seen here is due to changes in the ECM structure. Those analyses will be also necessary to clinically confirm our *in vitro* mechanistic observations, in which radiation generally decreased ECM fibre numbers and size in tumour cells.

## Conclusion

We characterised for the first time the ECM produced by MIBC cells surviving fractionated irradiation. Our data showed radiation mostly impacts the levels of non-structural ECM proteins (e.g. cytokines, proteases). At the structural level, glycoprotein levels were the most affected. We also showed radiation can disorganise the ECM structure, destroying and shortening ECM fibres. We found heterogeneity between cell lines, likely affected by different genomic backgrounds, and future research should explore whether COL fibrogenesis after irradiation depends on specific genomic mutations. We also showed that the ECM proteins identified as affected by irradiation were prognostic when assessed at the RNA level in untreated diagnostic biopsies of MIBC. Future research should explore changes in samples taken from tumours that recurred following radiotherapy to identify whether it might identify more radiation-specific effects. Last, we found that hypoxia-modifying treatment abrogated the prognostic capacity of the ECM genes studied suggesting further research should study the effect of CON on radiation-induced changes in the ECM produced by cancer cells.

## Acknowledgements

The work was funded by the NIHR Manchester Biomedical Research Centre (NIHR129943). The work was also supported by Cancer Research UK Major Centre (C147/A25254) and project grant (C1098/A9437; C2094/A11365; C13329/A21671) funding. The support of D. Knight, E. Keevill and J.N. Selley at the Bio-MS mass spectrometry core facility (RRID: SCR_020987) in the Faculty of Biology, Medicine and Health at the University of Manchester is gratefully acknowledged. The mass spectrometers used in this study were purchased with grants from BBSRC, Wellcome Trust and the University of Manchester Strategic Fund.

Work was carried out at The University of Manchester, Cancer Research UK – Manchester Institute, and the Manchester Cancer Research Centre, which provided infrastructure support and access to core facilities for the experiments performed in this study.

## Author Contributions

Conceptualisation: C.G.Q., S.F., A.C., R. R., C.M.W., V.B.; Methodology: C.G.Q., S.F., M.B., J.G.A., R. R.; Software: C.G.Q., V.B.; Validation: C.G.Q.; Formal analysis: C.G.Q., S.F., M.B., J.G.A., A.B., R.R., V.B.; Investigation: C.G.Q., S.F., M.B., J.G.A., E.P., R.R., A.B., Resources: P.H., N.D.J., E.H., R.A.H., N.P., A.C.; Data curation: C.G.Q., P.H., N.D.J., E.H., R.A.H., N.P., A.C; Visualisation: C.G.Q., A.B.; Supervision: R.R., P.H., A.C., C.M.W.; Project administration: R.R., K.R.; Funding acquisition: K.R., A.C., C.M.W.; Writing (original draft): C.G.Q.; Writing (review & editing): V.B., A.C., C.M.W.

## Declaration of interests

The authors declare that they have no conflict of interest.

## Supplementary figures

**Figure S1:**
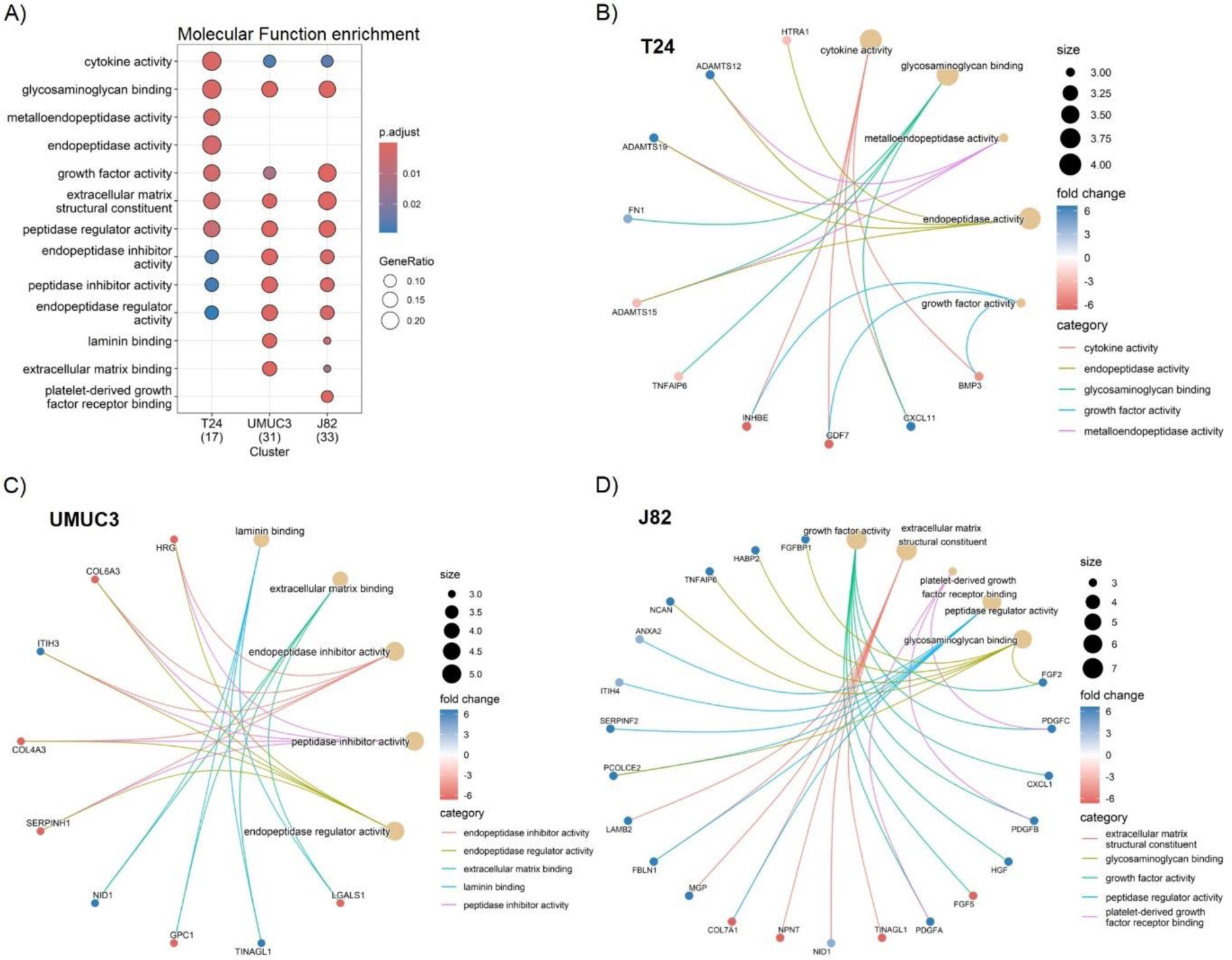
Fractionated radiotherapy partially mimicking SOC (27.5Gy, 2.75Gy daily over 2 weeks) consistently alters cytokines, glycosaminoglycan binding molecules, growth factors, peptidases and peptidase-regulators expression across all cell lines. Comparative Molecular Function pathway enrichment analysis (A) of significantly up (fold change >2, p.adj. <0.05) and downregulated (fold change <-2, p.adj. <0.05) ECM proteins for each individual cell line (T24, UMUC3, J82). Cnetplots show specific associations among significantly up and downregulated proteins and their corresponding enrichment terms for T24 (B), UMUC3 (C) and J82 (C). Ratio (A) represents the % of total proteins associated with each term (0 – 1 scale). A total of n=3 biological repeats were analysed per cell line.

**Figure S2:**
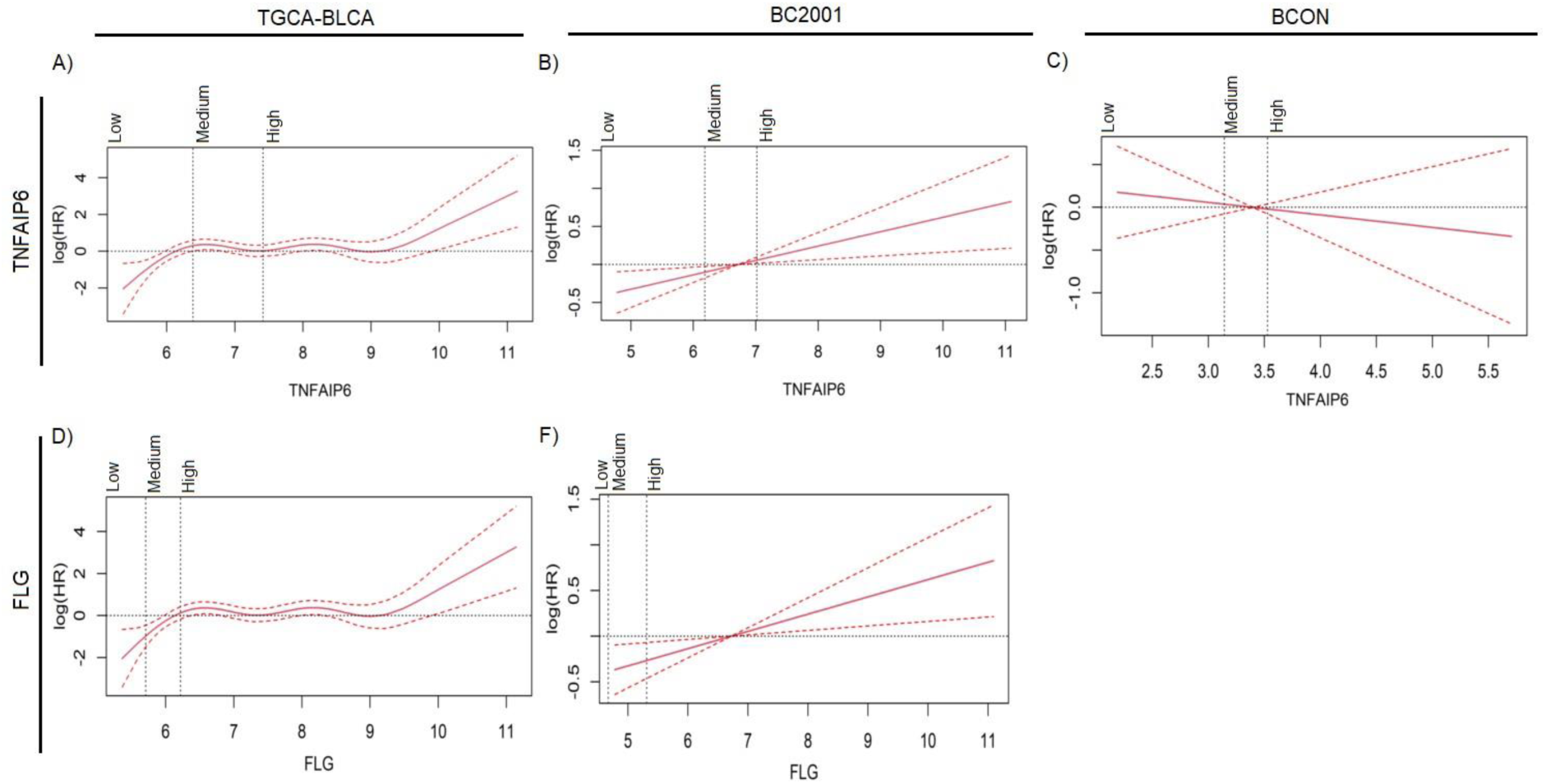
High *TNFAIP6 and FLG* mRNA expression levels are associated with a linear increase in mortality risk in muscle-invasive bladder cancer (MIBC). *TNFAIP6* (A-C) and *FLG* (D,M) expression was retrospectively validated in one cystectomy (TCGA-BLCA, n=397) and two radiotherapy (BCON [n=151]; BC2001 [n=313]) cohorts. Cut-off for patients with “High” (>67%), “Medium” (33 – 67%) or “Low” (<33%) hazard risk is based on each cohort’s tertiles median gene expression for *TNFAIP6* (J-L) and *FLG* (M,N) expression. No *FLG* expression data was available for the BCON cohort.

**Figure S3:**
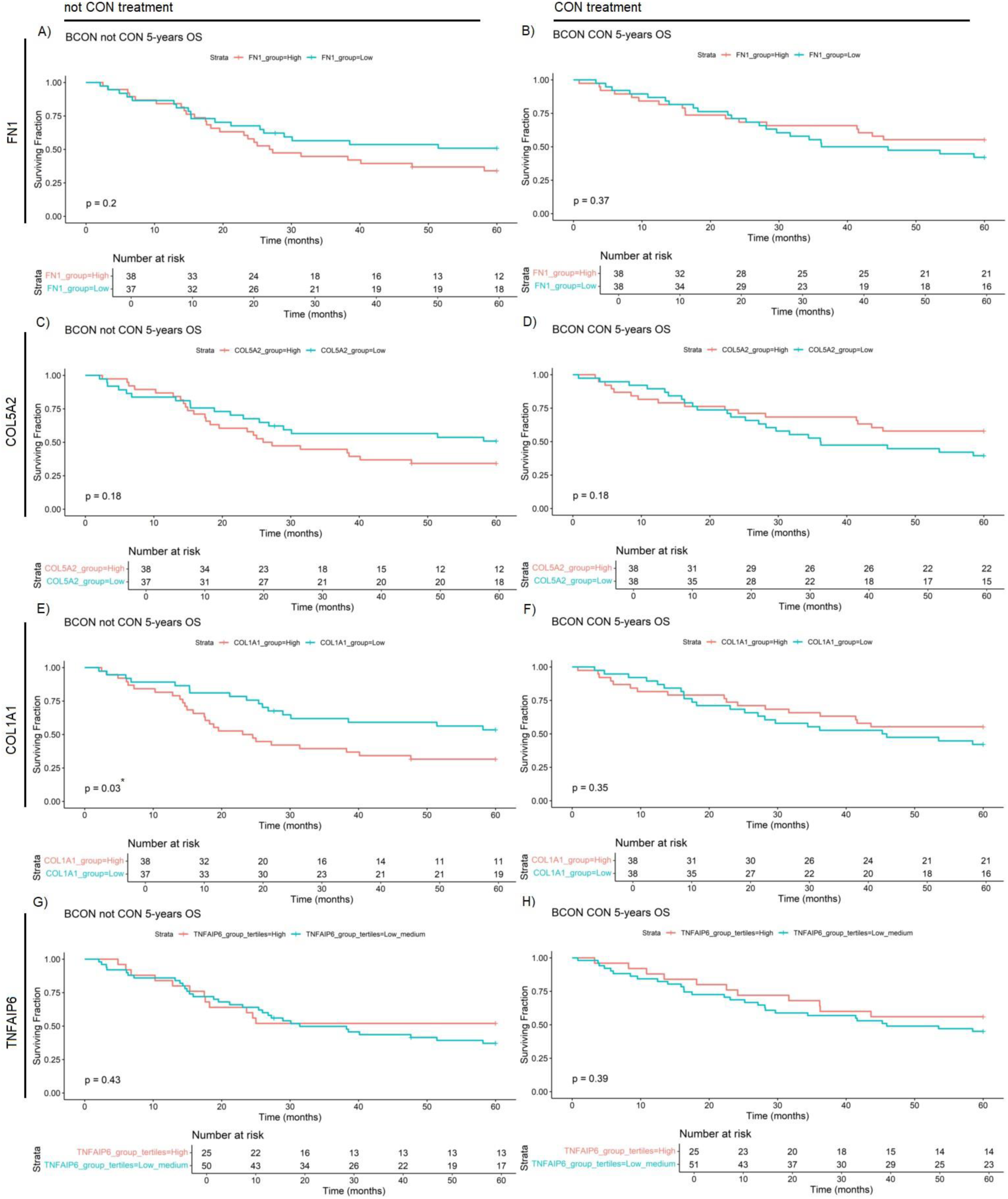
*COL1A1* mRNA expression level is a general prognostic marker for in muscle-invasive bladder cancer (MIBC) patients without carbogen and nicotinamide (CON) hypoxia-modifying treatment. *FN1* (A,B), *COL5A2* (C,D), *COL1A1* (E,F) and *TNFAIP6* (G,H) expression was retrospectively validated in the BCON cohort (n=151) after a split into the two CON treatment arms (n=75 not CON treatment; n=76 CON treatment). COL1A1 expression had significant prognostic value in patients without CON treatment (radiotherapy only) (E). No other gene had significant prognostic value, in either of the CON treatment arms of the BCON cohort. Patients were classified into “High” and “Low” based on each cohort’s median gene expression levels for *FN1* (A,B), *COL5A2* (C,D) and *COL1A1* (E,F) analyses. Patients were classified into “High”, “Medium” or “Low” based on each cohort’s tertiles median gene expression for *TNFAIP6* (G, H) expression. Significance was defined as p≤0.05, with * for p≤0.05, ** for p≤0.01, *** for p≤0.001 and **** for p≤0.0001.

**Figure S4:**
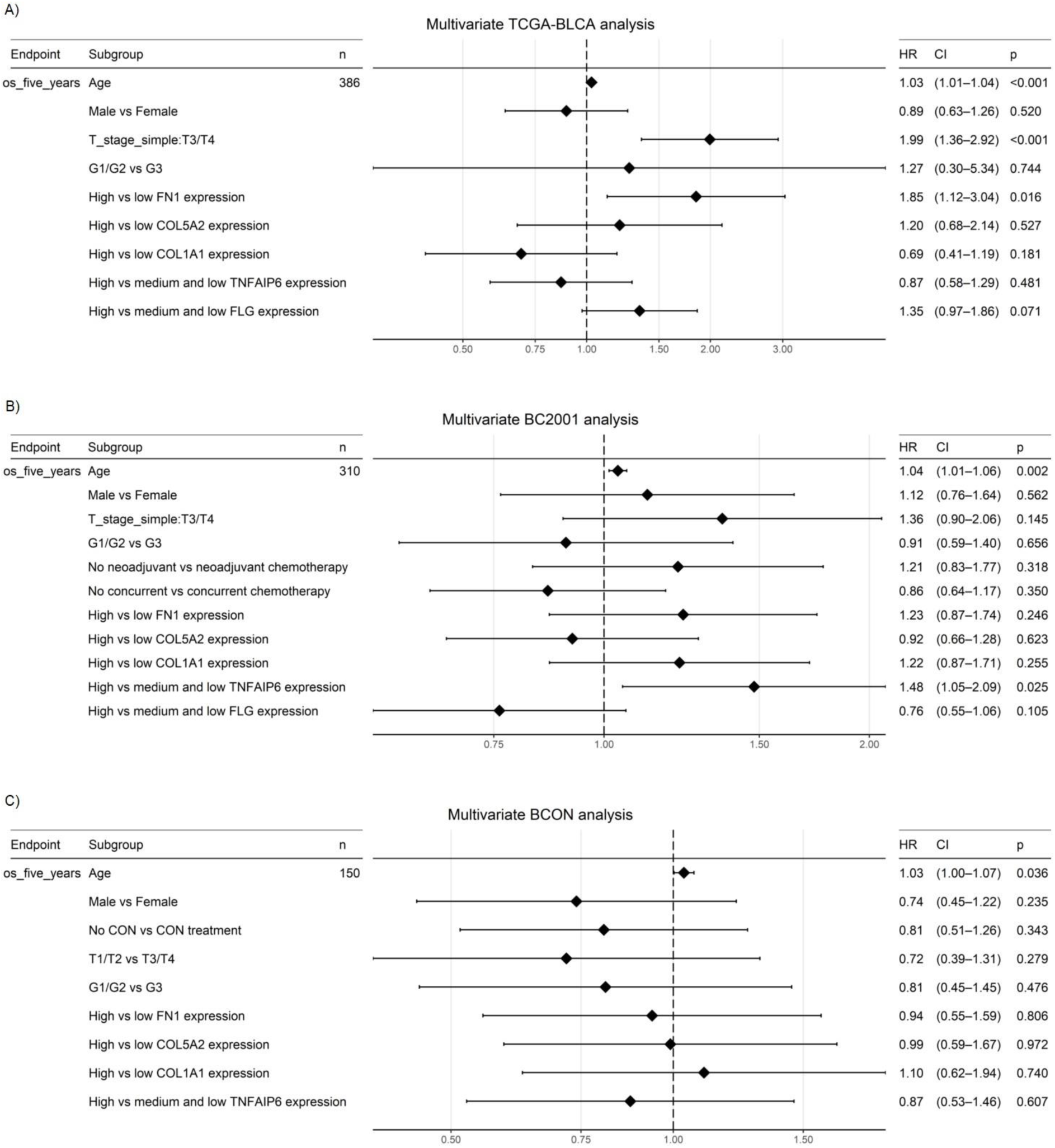
*FN1* and *TNFAIP6* are respectively independent prognostic markers in a cystectomy (TCGA-BLCA) and radiotherapy (BC2001) muscle-invasive bladder cancer (MIBC) cohort. Multivariate analyses show a 85% increase in mortality risk (p=0.016) was found for patients with “High” vs “Low” *FN1* expression in the TCGA-BLCA cohort (A). Furthermore, a 48% increase in mortality risk (p=0.025) was found for patients with “High” vs “Medium and Low” *TNFAIP6* expression in the BC2001 cohort (B). None of the remaining genes (*COL5A2*, *COL1A1*, *FLG*) showed significant value as independent prognostic markers. In addition, no gene was validated as an independent prognostic marker in the BCON cohort (C). No *FLG* expression data was available for the BCON cohort. Age was the only independent prognostic marker in all three cohorts (TCGA-BLCA, BC2001, BCON). T-stage was significant only in the TCGA-BLCA cohort (A). No other clinical variables or treatments significantly affected mortality risk. Patients were classified into “High” (≥50%) and “Low” (<50%) based on each cohort’s median gene expression levels for *FN1*, *COL5A2* and *COL1A1*. Patients were classified into “High” (>67%), “Medium” (33 – 67%) or “Low” (<33%) based on each cohort’s tertiles median gene expression for *TNFAIP6* and *FLG* expression. Significance was defined as p≤0.05, with * for p≤0.05, ** for p≤0.01, *** for p≤0.001 and **** for p≤0.0001.

## Notes

### Competing Interest Statement

The authors have declared no competing interest.

